# Transcontinental patterns in floral pigment frequencies among animal-pollinated species

**DOI:** 10.1101/2024.12.26.630375

**Authors:** Eduardo Narbona, Jose C. Del Valle, Justen B. Whittall, Melissa León-Osper, M. Luisa Buide, Iñigo Pulgar, Maria Gabriela Gutierrez Camargo, Leonor Patricia Cerdeira Morellato, Nancy Rodríguez-Castañeda, Victor Rossi, Katie Conrad, Joey Hernandez-Mena, Pedro L. Ortiz, Montserrat Arista

## Abstract

Flower coloration arises primarily from pigments that serve dual functions: attracting pollinators and mitigating environmental stresses. Among major flower pigment groups, anthocyanins and UV-absorbing phenylpropanoids are particularly notable for fulfilling both roles. Their importance suggests that both pigment types should be widespread in flowers. Here, we analyze the UV-Vis absorption profiles of major floral pigments to assess their potential protective role against UV radiation and demonstrate that this protection is largely confined to UV-absorbing phenylpropanoids. We also analyzed the floral pigment composition of 926 animal-pollinated species from California, southern Spain, and southeastern Brazil. UV-absorbing phenylpropanoids were ubiquitous, while anthocyanins occurred in ∼56% of species, carotenoids in ∼37%, and chlorophylls in ∼17%. Pigment frequencies varied with abiotic and biotic factors, such as light environment in Spain, and pollinator type, such as insect versus hummingbird, in California. Despite these local differences, our findings reveal a consistent regional distribution of floral pigments.

## Introduction

The vast majority of flowering plants depend on pollinators for reproduction (∼90%, ^1^). Plants produce different types of floral signals to attract pollinators and ensure seed production, with flower color being one of the most important visual cues perceived by pollinators^2^. Since color signals received by pollinators are mainly determined by floral^3,4^, and pollinators are important selective agents^5^, we can expect that floral pigment composition is a target of selection^6^. Nonetheless, the specific role of pigments in pollinator adaptation remains unclear, as does the extent to which other selective forces act on floral pigment composition^7,8^.

A growing body of evidence suggests that floral pigments may also be crucial for stress protection^9–11^. All major types of pigments, such as phenylpropanoids — including UV-absorbing phenylpropanoids (UAPs hereafter; mainly hydroxycinnamic acids, flavonols, flavones) aurones, chalcones, and anthocyanins — as well as carotenoids, chlorophylls, and betalains, possess antioxidant properties to a greater or lesser extent^12–15^. However, UAPs and anthocyanins seem more effective in mitigating extreme temperatures, drought, pathogens, and herbivores, as demonstrated in vegetative tissues^16–17^. UAPs are recognized for their sunscreen properties due to their ability to absorb UV-A and UV-B radiation^18–19^. In vegetative organs, UAPs have played a critical role in protecting against UV radiation during the terrestrialization of plants^19–20^. However, their presence in flowers has received limited attention ^3,21^ and has been investigated only in specific lineages (e.g.^22,23^). In parallel, the frequency of anthocyanins at community or regional scales is frequently deduced based on floral coloration rather than through direct pigment analysis (e.g.^24–26^). The concurrent accumulation of anthocyanins and UAPs in flowers may not only enhance the antioxidant capacity of these pigment groups but also simultaneously support pollinator attraction ^13–27^. A critical challenge in this context is the assessment of the prevalence of UAPs and anthocyanins in the flowers of angiosperms.

Given that UV irradiance is a significant stressor for terrestrial plants^28^, and pigments likely play a role in helping flowers mitigate its effects^13,16^, here we first reviewed the UV-Vis absorption spectra of most important pigment groups present in flowers. We found that UAPs are the most common pigment with a considerable capacity to absorb light in the UV region. We then analysed flower pigments in 936 animal-pollinated species spanning 115 families and 486 genera from three geographic regions: California, southern Spain, and southeastern Brazil. Our main hypothesis is that if the accumulation of floral UAPs is advantageous for coping with a nearly universal environmental stress, such as UV radiation^20,28,29^ they should be more frequent than anthocyanins in angiosperm flowers.

Additionally, the accumulation of floral UAPs benefits pollinator attraction, as they are common components of UV floral guides^30,31^. On the other hand, since habitat influences light environment and pollinator color perception, Endler^32^ predicted that flowers in forest shade should be yellow or yellow-green to maximize brightness. Thus, given that these flower colors are a consequence of the presence of carotenoids, aurone-chalcones, or chlorophylls, we expect a high frequency of these pigments in flowers of shaded habitats. Finally, bird-pollinated flowers typically produce a red UV-absorbing coloration that attracts birds while avoiding bees^33–35^, leading to high accumulation of UAPs combined with anthocyanins and/or carotenoids^36–38^. Therefore, we predict a high frequency of these three pigments in hummingbird-pollinated flowers with respect to insect-pollinated flowers. We found that UAPs are ubiquitous independent of flower color and the presence of other floral pigments. Notably, while floral pigment composition was largely consistent across the three geographic regions studied, local variations were influenced by differences in pollination systems and light environments.

## Results and Discussion

### Absorption properties of major groups of floral pigments

We reviewed the literature to find out which part of the light spectrum is absorbed by the four main classes of pigments, namely phenylpropanoids, carotenoids, chlorophylls and betalains, and their derivatives more commonly found in flowers (Figure 1). Within the phenylpropanoids, hydroxycinnamates (aka hydroxycinnamic acids), flavones and flavonols exclusively absorb in the UV range, but hydroxycinnamates absorb mainly in the UV-B region of the spectrum (peaks ≈ 280 and 330 nm) and flavones and flavonols in the UV-A region (peaks ≈ 310 and 390 nm; ^39–41^). Aurones and chalcones (treated together herein) absorb in the UV-blue region whereas anthocyanins mainly absorb in the green-blue region^40–42^. Flowers may contain other groups of flavonoids such as flavanones, isoflavones, catechins or epicatechins, but they are relatively rare compared to the aforementioned groups^42,43^. Carotenoids absorb mainly in the blue-green spectral region of the visible light and chlorophylls absorb in the blue and red regions (although chlorophyll *a* has a relatively moderate absorption ability in the UV-A region) (Harborne, 1984; Ohmiya, 2011). Lastly, betalains show absorption spectra similar to anthocyanins^13,40^, but they are restricted to a few families within the order Caryophyllales^44^. In summary, the only pigments with a considerable capacity to absorb light in the UV region of the spectrum are the UAPs (i.e., hydroxycinnamates, flavonols and flavones) and aurones-chalcones. This would explain why in photosynthetic tissues UV-B exposure generally promotes the biosynthesis of hydroxycinnamates, while UV-A exposure stimulates the production of flavonols and flavones^18,45,46^ (see also^47^). Although evidence in flowers is more limited, available studies suggest a similar trend^11,48^.

**Fig. 1.**
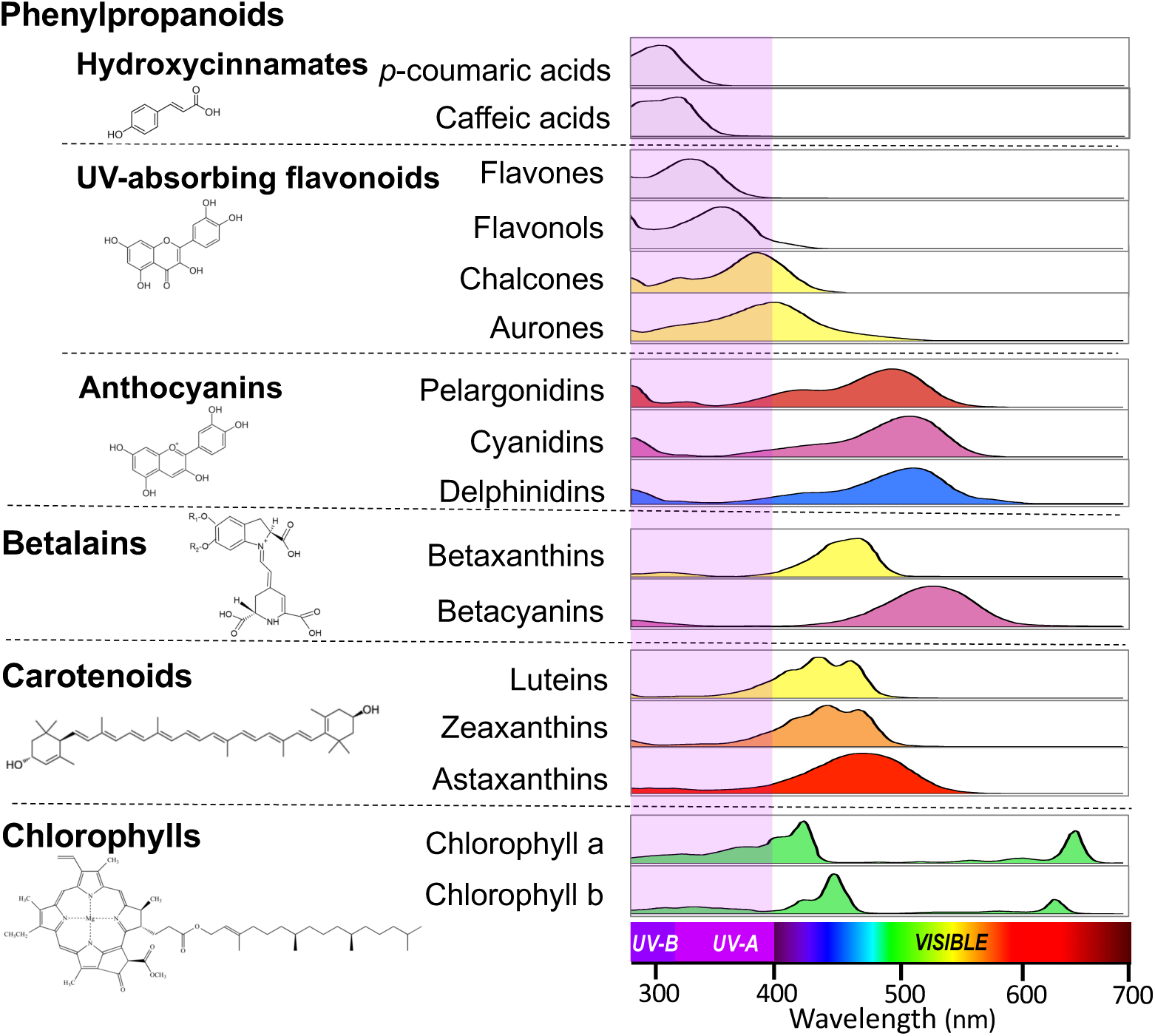
Ultraviolet (UV) and visible absorption spectra of major pigment groups contributing to flower coloration. Due to their structural diversity and distinct contributions to floral coloration, phenylpropanoids were categorized into hydroxycinnamates, UV-absorbing flavonoids, aurones-chalcones, and anthocyanins. The spectral region corresponding to each pigment type is color-coded based on the human perception of reflected light from petals. The violet-shaded area highlights the UV-A and UV-B regions of the light spectrum (400-315 nm and 315-280 nm, respectively) and shows that hydroxycinnamates and UV-absorbing flavonoids are the pigments with the highest light absorption capacity in this range. For each group of pigments, a normalized absorption spectrum of an example compound is shown (see Methods). The structures of *p*-coumaric acid (hydroxycinnamate), quercetin (UV-absorbing flavonoid), cyanidin (anthocyanin), betacyanin (betalain), lutein (carotenoid), and chlorophyll a (chlorophyll) are depicted.

### UV-absorbing phenylpropanoids are pervasive in flowers

We performed a biochemical analysis of flowers of 926 animal-pollinated species from diverse habitats, 442 in California, 381 in southern Spain, and 103 in south-eastern Brazil (Supplementary Data 1). Using a differential extraction method followed by an analysis of absorbance spectra (see details in Methods), we were able to identify six major groups of pigments: UAPs (hydroxycinnamic acids, flavones and flavonols), aurones-chalcones, anthocyanins, chlorophylls, carotenoids, and betalains (Figure 2A). Notably, the presence of UAPs was almost ubiquitous in species from the three regions (> 99.8%; Figure 2B). Only one species from California, *Geum macrophyllum* (Rosaceae), lacked UAPs in the flowers.

**Fig. 2.**
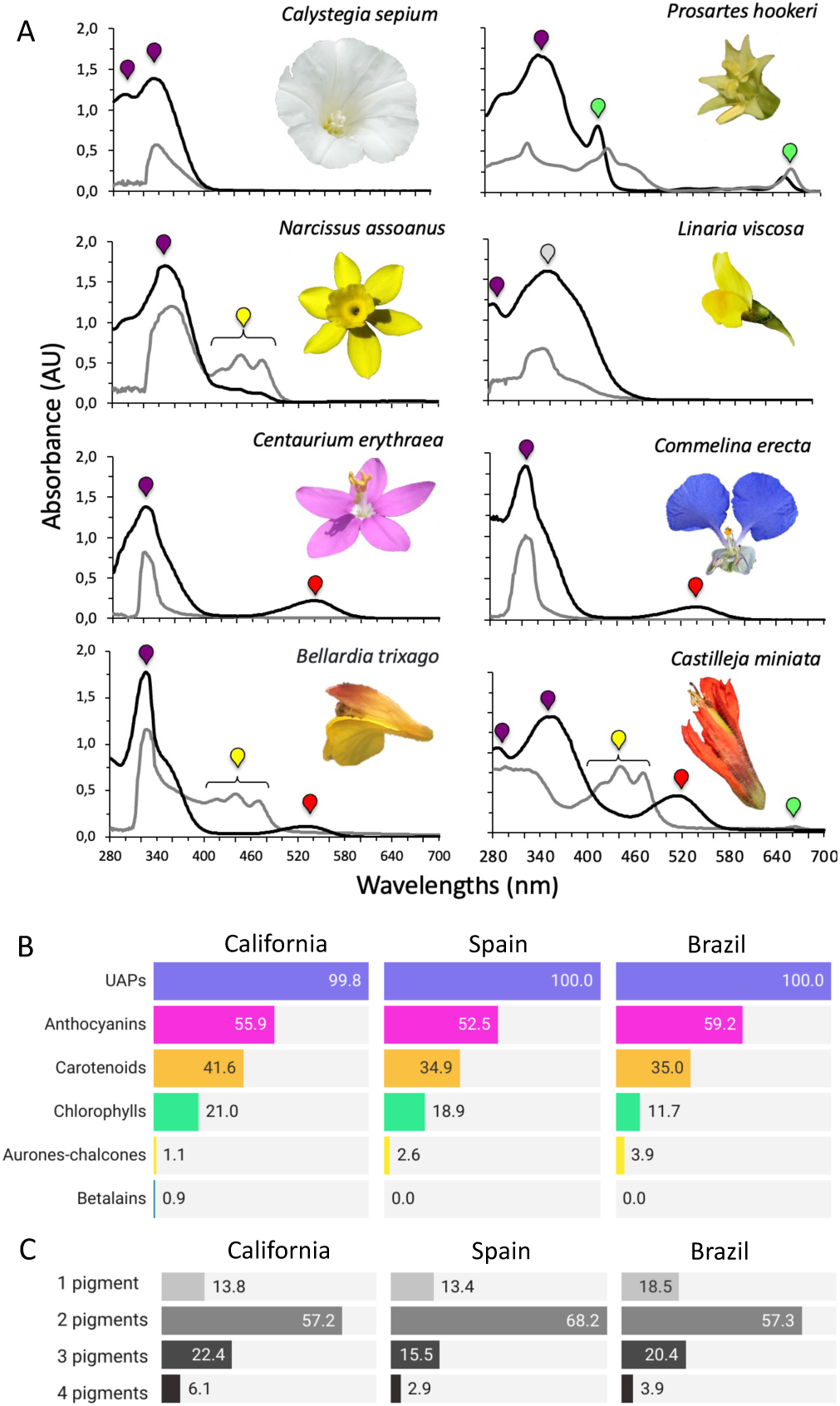
Pigment identification and frequency in the studied regions. **A** Examples of absorbance spectra for acidified methanol and acetone extracts (black and gray lines, respectively) from flowers of different colors, caused by a combination of pigment classes. The differential solvent extraction and the characteristic absorption peaks of each pigment type allow identification of the pigment types present in the crude extract. Colored drops are used to show the identifiable peaks of the pigment types. In the methanol extracts, violet drops are used for UV-absorbing phenylpropanoids (UAPs), grey drops for aurones-chalcones, red drops for anthocyanins, and green drops for chlorophylls. Betalains not included due to their rarity. In flowers that accumulate several kinds of pigments, some pigment peaks may overlap, but the characteristic types of each pigment may allow them to be differentiated (see methods). Acetone extracts are used to identify distinctive three-peaked carotenoids and are marked by a yellow drop. Betalains show a similar behavior to those of anthocyanins but are not represented because of their rarity. An example of a species showing a single pigment class (*Calystegia sepium*, Convolvulaceae, Spain; white flowers with UAPs). Examples of species with two pigments classes: *Prosartes hookeri* (Liliaceae, California; green flowers with UAPs and chlorophylls), *Narcissus assoanus* (Amaryllidaceae, Spain; yellow flowers with UAPs and carotenoids), *Linaria viscosa* (Plantaginaceae, Spain; yellow flowers with UAPs and aurones-chacoles), *Centaurium erythraea* (Gentianaceae, Spain; pink flowers with UAPs and anthocyanins), *Commelina erecta* (Commelinaceae, Brazil; blue flowers with UAPs and anthocyanins). An example of species with three pigments (*Bellardia trixago*, Orobanchaceae, Spain; yellow-red flowers with UAPs, anthocyanins and carotenoids), and with four pigments (*Castilleja miniata*, Orobanchaceae, California; red flowers with UAPs, anthocyanins, chlorophylls and carotenoids). **B** Percentage of species containing each type of floral pigment in California, S Spain, and SE Brazil (N = 442, 381 and 103, respectively). Note that in a flower each pigment type may appear alone or coexist with other pigment types. **C** Frequency of the number of floral pigment types present in species from California, S Spain, and SE Brazil (N = 442, 381 and 103, respectively).

Anthocyanins were the second most abundant pigment group, present slightly more than half of the species in the three study areas (mean 55.9%), while carotenoids appeared in about one-third of the species (mean 37.2%). Chlorophylls ranged from 12% to 21% (mean 17.2%), while aurone-chalcone pigments were infrequent, found in less than 4% of species. We also confirm that betalains are uncommon pigments, present in only four sampled species of California (0.9%; Figure 2B). Using ultraviolet microscopy and chromatography on petals of 201 species, Kay et al.^3^ found UV-absorbing flavonoids and anthocyanins in half of the species. Recent studies on Brassicaceae, Orchidaceae, and Solanaceae suggest that UAPs accumulate in flowers regardless of color or the presence of floral guides^22,23,49,50^. Our results clearly show that the presence of UAPs in flowers is pervasive in the species studied and, presumably, in all angiosperms. This finding aligns with the ubiquity of UAPs in vegetative tissues, where they serve multiple protective functions^16,19^. For instance, floral UAPs may counteract oxidative damage induced by UV radiation or drought in petals^51,52^ or help to maintain cellular turgor through sugar signaling^53^.

Although we found that flowers can accumulate up to four types of pigment groups, most species presented only two types, with 57% of species sampled in California and Brazil and 68% of species sampled in Spain falling in this category (Figure 2C and S2). Because of the omnipresence of UAPs in flowers, the most common combination of two pigments was UAPs + anthocyanins (58-66%; Figure S3). In petal cells, anthocyanins typically accumulate in vacuoles^54^, where they can be found alone or bonded to UAPs by copigmentation or other molecular interactions to stabilize or intensify the color^42,55^. This possibility of association with UAPs and the great diversity of anthocyanin derivatives is proposed to generate the enormous range of colors and color pattern of angiosperm flowers^42,43,56^. In addition to the effect on floral color perceptible to humans and pollinators, the association between UAPs and anthocyanins results in anthocyanins gaining a substantial light absorption capacity in the UV-A and/or UV-B region^18,57,58^. Thus, accumulation of UAPs and anthocyanins in flowers may be advantageous to the plant and can be shaped by both biotic and abiotic factors^13,59^.

### UV-absorbing phenylpropanoids are typically distributed throughout the entire petals

Most species accumulated UAPs in the entire petals (80.7%, 86.9% and 92.3% in California, S Spain and SE Brazil, respectively) and the rest of species had UV–patterned flowers including bullseyes, veins, rays, spots, and/or differently colored petals/tepals (e.g. flowers of Fabaceae or Orchidaceae, Figure 3, Supplementary Data 1; see also^60^). In UV photography, a few flowers appeared mostly UV-reflective across the corolla (see other examples in^31,61^), but in our absorbance analysis, these flowers were positive in UAPs. This apparent contradiction may be due to the greater accuracy of the absorbance analysis, which detected UAPs even at low concentrations, or to the possibility that UAPs accumulate on the outer sides of petals^62^. Notably, UAPs may protect floral tissues even when confined to floral guides. For instance, UAPs accumulation in bullseyes may protect reproductive structures from UV radiation or enhance petal resistance to desiccation^11,63,64^.

**Fig. 3.**
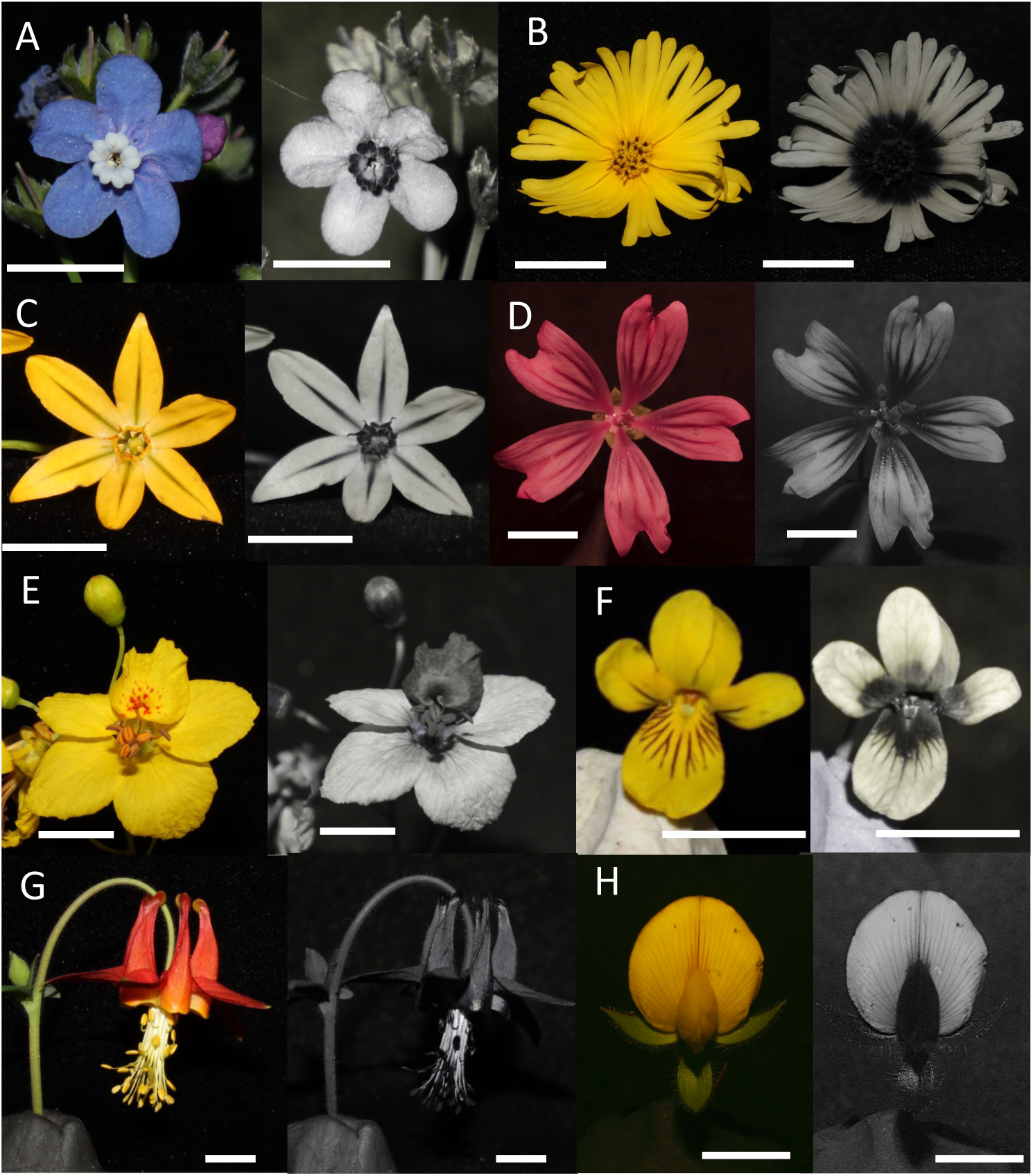
UAPs location in entire petals or in UV spatial color patterns. Visible (left) and UV (right) photography allow to find UV-absorbing areas in the corolla caused by the accumulation of UAPs. Scale bars = 10 mm. **A** *Adelinia grandis* (Boraginaceae, California) showing a UV-absorbing bullseye in the corolla base. **B** *Madia elegans* (Asteraceae, California) showing a highly UV-absorbing bullseye in disk florets and base of ray florets. **C** *Triteleia ixioides* (Asparagaceae, California) showing UV-absorbing rays on the tepals and corona, in this case is accompanied by chlorophylls causing green rays in visible photography. **D** *Malva sylvestris* (Malvaceae, Spain) showing UV-absorbing veins and bullseye, in this case is accompanied by high concentration of anthocyanins causing purple rays in visible photography. **E** *Parkinsonia florida* (Fabaceae, California) showing visible dots and one petal with high UV absorption. **F** *Viola glabella* (Violaceae, California) showing UV and visible veins and UV-absorbing bullseye. **G** *Aquilegia formosa* (Rannunculaceae, California) showing homogeneous UV-absorbing corolla. **H** *Ononis pubescens* (Fabaceae, Spain) showing visible and UV veins, and keel and wings with high UV absorption.

### Flower pigment frequencies are consistent across regions

We observed striking similarities in floral pigment frequencies across the three regions analyzed (Figure 2B and S1). A survey of British flora, which categorized pigments into broad color groups—such as pink, red, blue, violet, and purple (anthocyanins) and yellow (carotenoids)—similarly reported a higher prevalence of anthocyanins compared to carotenoids^24^. However, the anthocyanin frequency observed in their study was lower than in ours. This discrepancy likely stems from their method, which considered only the predominant pigment responsible for the main flower color, whereas our approach accounted for all pigments present in the flowers. Using the same flower color categorization as Warren and MacKenzie^24^, a higher frequency of anthocyanins compared to carotenoids has been consistently observed across various regions, including France, Scandinavia, Canada, tropical Africa, Australia, and Java^65,66^, as well as on a global scale^67^.

Although the consistent frequencies observed across regions worldwide are likely driven by multiple factors, phylogenetic similarity can be ruled out given the distinct floras of these regions. In fact, the frequency of shared families in our study was also low: 30.6% (38 of 124 families) between California and Spain, 15.9% (14 of 88) between Spain and Brazil, 14.0% (14 of 100) between California and Brazil, and only 7.6% (12 of 157) among the three regions. Both random genetic drift and deterministic processes, including biotic and abiotic selective factors, may influence the assembly of flower colors within populations^68–70^, ultimately shaping the distribution of flower pigment groups in communities^26,71^. Pollinators are considered important selective agents on flower color, and are involved in flower color convergence, facilitation, character displacement and competitive exclusion processes, which may originate flower color clustering or overdispersion^71–73^. All these deterministic processes assume that pollinators have visual preferences for certain flower colors^33,74^.

Hymenopterans show differences in naive preferences among species and have complex learning mechanics^75^, but in general they prefer blue-violet and yellow over other flower colors, with a higher bias towards blue-violet^76^. Thus, the higher frequency of anthocyanins compared to carotenoids observed in our three study regions may be explained by the floral color preferences of hymenopterans, which are the most frequent pollinators in each region. Although less studied, herbivory—another biotic factor—may also influence flower color within communities, with reported cases suggesting a selective advantage for anthocyanins over carotenoids^77,78^. Lastly, biochemical and molecular factors may contribute to the higher prevalence of anthocyanins compared to carotenoids. For example, it is well-documented that abiotic factors such as temperature, precipitation, and solar radiation can increase the frequency of species that accumulate floral anthocyanins or UAPs^25,29,79^. In addition, petal anthocyanins may be less costly to produce and store, as they accumulate in vacuoles, whereas carotenoids are stored in more complex chromoplasts^56,80^. Anthocyanins and the ubiquitous UV-absorbing phenylpropanoids (UAPs) share the first half of their biochemical pathways, which may facilitate the production of floral anthocyanins^17^ and, ultimately, the diversification of angiosperms, enabling them to produce colorful flowers that attract pollinators^13–20^.

### Pigment composition is influenced by both abiotic and biotic factors

We tested whether the overall frequency of major floral pigment groups is influenced by different light environments—shaded (forest, riparian) vs. exposed (grasslands, coastal, rocky, etc.)—in California and S Spain. In both regions, most of sampled species belong to the exposed habitats (82% and 88.8%, respectively). We found a trend toward a lower frequency of anthocyanins and carotenoids and a higher frequency of chlorophylls in shaded environments compared to exposed environments, but the differences were only significant for chlorophylls in Spain (Figure 4 and Table S1). These findings partially support Endler’s prediction^32^ of a higher prevalence of yellow-green flowers in shaded environments and highlight the potential role of floral chlorophylls in attracting insect pollinators^81,82^.

**Fig. 4.**
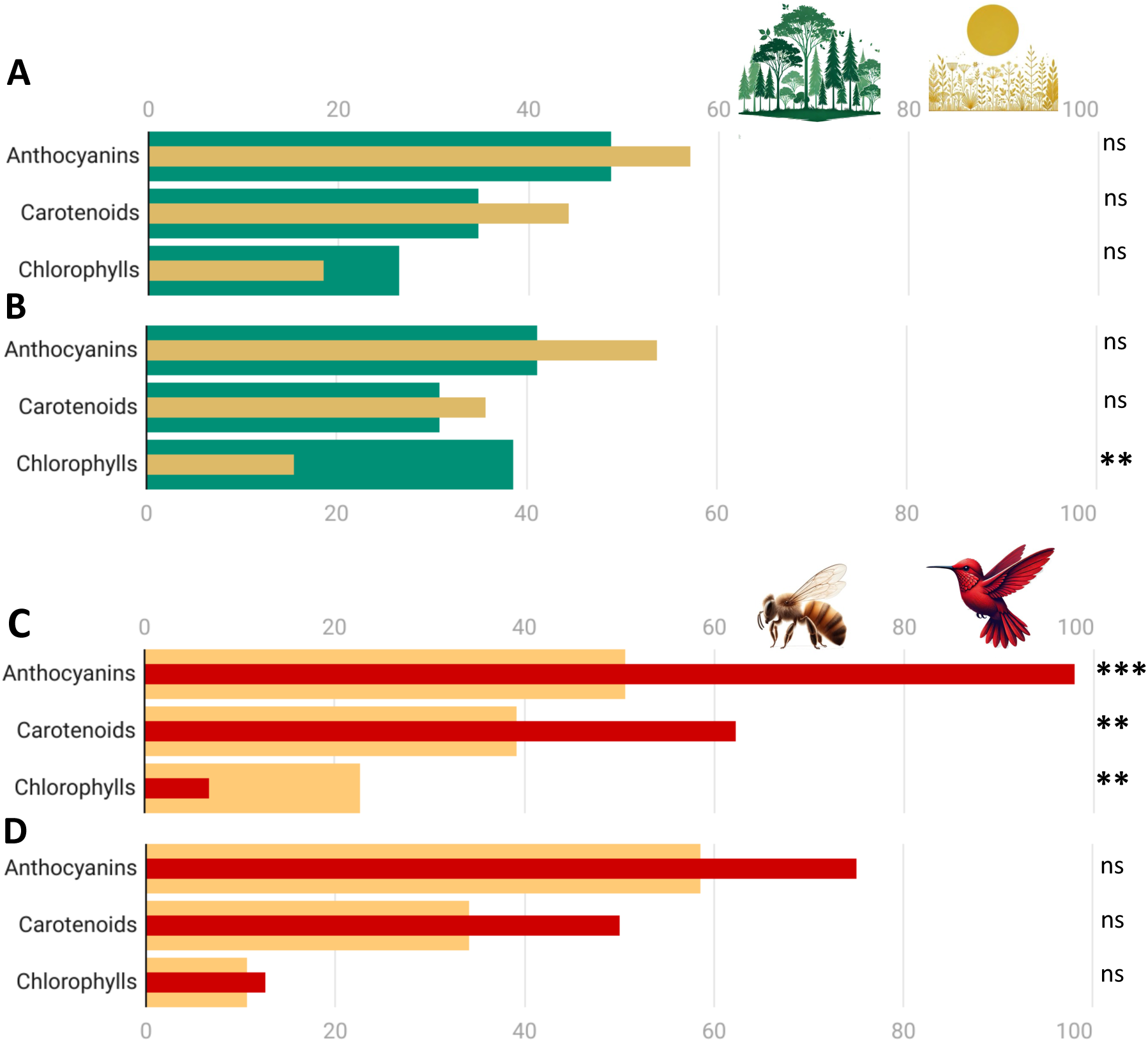
Variation in the frequency of floral pigment types across different light environments and pollination systems. **A** Comparison of the frequencies of the three main pigment groups between exposed (light brown) and shaded (green) light environments in California (N = 328 and 72, respectively). **B** Comparison of the frequencies of the three main pigment groups between exposed (light brown) and shaded (green) light environments in S Spain (N = 309 and 39). **C** Comparison of the frequencies of the three main pigment groups between insect (light orange) and hummingbird (red) pollination systems in California (N = 393 and 45, respectively). **D** Comparison of the frequencies of the three main pigment groups between insect (light orange) and hummingbird (red) pollination systems in SE Brazil (N = 94 and 8). Symbols on the right show the results of permutation tests to compare the frequency of each pigment group between light environments and pollination systems (see statistical results in Supplementary Table 1 and 2). n.s. not significant, *p < 0.05, **p < 0.01, ***p < 0.001. Images of light environments and pollination systems were generated by AI image generator DALL-E.

We also examined if insect- and hummingbird-pollinated species differed of floral pigment distribution. Our database includes 9.7% of species categorized as hummingbird-pollinated in California and 12.8% in southeastern SE Brazil, which aligns with previous reports from both regions^34,83,84^. In California, the frequency of anthocyanins and carotenoids in bird-pollinated flowers was nearly double that those of insect-pollinated flowers, while chlorophyll showed the opposite trend (Figure 4 and Table S2). The red color is a defining characteristic of the hummingbird pollination syndrome^85^ (Pauw, 2019). This coloration arises from the accumulation of red anthocyanins or a combination of blue-pink anthocyanins and yellow carotenoids^36,38^. Several studies have demonstrated hummingbirds’ preference for these pigments^38,85^. Our findings confirm the prevalence of these pigments across 16 plant families in California (Supplementary Data 1) and suggest that hummingbirds are driving the evolution of floral pigmentation in this region. A higher frequency of anthocyanins and carotenoids was observed in hummingbird-pollinated flowers form SE Brazil, but differences were not statistically significant. This may be due to the relatively high prevalence of hummingbird-pollinated species with yellow or white flowers in the Brazilian cerrado, likely reflecting a less specialized hummingbird pollination syndrome^34,84^.

### Conclusions

Our trans-continental floral biochemical results highlight the ubiquitous nature of UAPs compounds in animal-pollinated flowers. We argue that UAPs may have a dual role in flowers, attracting pollinators and protecting against environmental stresses^13^. The genetic architecture for UAP biosynthesis is present in early land plants, allowing them to cope with UV radiation and thermal stress^20,28^. The main (ancestral) function of UAPs in floral tissues could be to protect cells from environmental stressors and the role of pollinator attraction could then be an exaptation^86^. The omnipresence of floral UAPs may be explained by their origin during the terrestrialization of plants, having been retained in all plants due to phylogenetic inertia and favored due to their versatility as environmental protectors^87^.

We found that floral pigment frequencies are consistent across the three studied regions, a pattern that likely extends globally. Our results offer a more comprehensive understanding of floral pigment abundance and further strengthen previous studies based on flower color or theoretical models^25,33,70^. The causes of these stable distributions are not well understood but likely reflect the flower color preferences of insects—the most frequent pollinators—as well as underlying biochemical and molecular factors, warranting further investigation. The variation in pigment frequency between exposed and shaded environments, as well as between insect- and hummingbird-pollinated flowers, suggests that eco-evolutionary processes are acting at local scales^7,25,88^.

## Methods

### UV-Vis absorption spectra of main flower pigment groups

We reviewed references presenting spectral data of pigments extracted in methanol, as it is one of the most common solvents used for plant pigments^30,89^. For each main pigment group, we selected the most representative or frequent subgroups found in flowers^42,43,56,90,91^ and present a normalized absorption spectrum of an example compound in Figure 1. The compounds used are show in Supplementary Table 3. Although there may be variation in the shape and wavelength at which the peak of maximum reflectance occurs between different pigment types within the same group, this variation is usually quite small compared to the differences between pigment groups^30,40,90^.

### Sampling for the floral pigment composition analyses

In California (California Floristic Province), we collected flowers from a total of 442 native species belonging to 249 genera and 69 families, and in southern Spain, we collected a total of 381 native species belonging to 198 genera and 56 families (Supplementary Data 1). In southeastern Brazil (National Park of Serra do Cipó, Minas Gerais, Brazil), we collected flowers from 103 species belonging to 72 genera and 32 families. Samples were collected from 2019 to 2023 in California and Spain and in 2019 in Brazil.

Both California and S Spain have a Mediterranean climate with typically hot, dry summers and cool, wet winters^92^. In SE Brazil (Minas Gerais state), the climate is tropical, with a cool dry season, a warm wet season, and frequent fires and strong winds during the dry-to-wet transition^93^. In California and S Spain, we sampled broadly across habitats by collecting in grasslands, shrublands, forests, riparian, rocky, wetlands, coastal, mountain and deserts communities throughout the year. In SE Brazil, we surveyed the rocky grasslands at high altitudes (>900 m a.s.l., Campo rupestre *sensu stricto*) and the woody savanna vegetation (Cerrado *sensu stricto*). The main pollinators in the three study regions were insects (primarily hymenopterans, but also dipterans, lepidopterans, coleopterans, among others; Moldenke 1976; Reverté *et al*., 2016; Camargo *et al*., 2019). However, in both California and SE Brazil, hummingbirds serve as efficient pollinators, with approximately 5– 12% of the flora exhibiting the typical "hummingbird-pollination syndrome"^34,83,94^. We analyzed the pigment content of floral parts with the highest advertisement display, typically petals or tepals, at the anthesis stage.

In all cases, we picked flowers or inflorescences at the anthesis stage of several individuals to take into consideration possible inter-individual variation in pigment composition. Additionally, 56 species were collected in two different populations to test the possibility of pigment variation between populations; in all cases, the qualitative pigment profile was identical and, thus, we used the data of the first population analyzed. Voucher specimens of species were deposited in the herbaria of University of Seville (Herbario SEV), University Pablo de Olavide (UPOS), Instituto de Biociências at the Universidade Estadual Paulista at Rio Claro (HRCB), and in JBW’s personal collection at Santa Clara University.

We sampled “attraction units”^95^, which in most species coincide with individual flowers but in some species, we considered the entire inflorescence (e.g., spadix of Araceae spp. or compound inflorescence of Asteraceae spp.). We analyzed pigment content of the floral piece that generates the highest advertising display, usually petals or tepals, but floral bracts were also examined in some species (e.g, Aristolochiaceae spp., Euphorbiaceae spp., Castilleja spp., or *Rhynchospora* spp.).

### Extraction, identification, and quantification of major pigment groups

Pigment extraction was carried out the same day or the next day after collection, keeping the flowers refrigerated at 4 °C until pigment extraction. We used two solvents to extract and separate the principal pigment classes in each sample: methanol with 1% HCl and pure acetone^40,96^. Methanol solution is particularly effective in extracting UAPs, anthocyanins, betalains, and chlorophylls, whereas acetone mainly extracts carotenoids and chlorophylls^40,89,97^. We used methanol as a solvent instead of ethanol or aqueous methanol solutions due to its superior efficiency to extract polar compounds. Acidification of methanol with HCl contributes to the stabilization of anthocyanins^97^. We placed approximately the same amount of floral tissue (5-25 mg of fresh weight) in two microtubes containing 1.5 ml of each solvent, which was kept in the freezer at -20 °C until further analysis. In California, we used 5-10 silica beads for tissue homogenization for 2-3 minutes followed by 5 minutes of centrifugation at maximum speed. In Spain and Brazil, homogenization was employed solely in cases involving thick tepals or floral bracts^99^.

We obtained the absorbance spectra of the samples in each solvent by means of two ultraviolet-visible (UV-vis) spectrophotometers. We used a Multiskan GO microplate spectrophotometer (Thermo Fisher Scientific Inc., MA, USA) with acetone-compatible polypropylene 96-well microplates and a Drawell double beam UV/Vis spectrophotometer (DU-8800DS model) with 1-cm quartz cuvettes, respectively. Previous studies showed that pigment quantification using these two types of containers provides nearly identical results^100^. We set the scan mode from 280 to 700 nm with 1 nm steps at a constant temperature of 22 ° C. We selected wavelengths above 280 nm because below that all phenolic compounds are characterized by a UV band II that peaks at 240-275 nm making it useless for differentiation^39,40^. We used 150 μl and 500 μl per sample in 96-well microplates and 1-cm quartz cuvettes, respectively. Following the technical specifications of the spectrophotometers, concentrated pigment extracts were diluted to obtain absorbance values below 2.0 absorbance units (AU) to guarantee reliable measurements.

Since our objective was to identify major pigment classes, we performed spectrophotometric analyses to obtain the absorption spectra of the sample extracts. Although with a lower within-pigment classes identification resolution, spectrophotometric analyses represent a reliable, fast, and inexpensive alternative to HPLC separation with mass or NMR spectrometry^97,101^. The fact that the major pigment classes have a particular light absorption spectrum with distinguishable peaks^39,102^, allowed us to identify the presence of such pigments in the floral extracts^27,40,102^. Thus, carotenoids show three distinctive peaks between 400 and 500 nm with a major peak around 450 nm, whereas chlorophylls show two main peaks at 415-460 nm and 650-665 nm^102^. All floral samples with the presence of chlorophylls showed a peak around 418 nm, suggesting that the most abundant is the chlorophyll a type^102^. We distinguish three major groups of phenylpropanoids: anthocyanins, aurones-chalcones, and UV-absorbing phenylpropanoids (UAPs). The flavonoid anthocyanins show a characteristic peak around 475-560 nm, whereas those of aurones-chalcones peak at 350-430 nm^40,42^. UAPs include the groups of flavonoids flavanones, flavones, and flavonols showing principal peaks from 280 to 360 nm, and some groups of hydroxycinnamates, such as cinnamic, caffeic, ferulic, p-coumaric, and sinapic acids with a peak at 280-330 nm^39,40,41,89^. The different groups of UAPs show distinctive peaks, but most of them overlap, which precludes their differentiation using this methodology^103^. Spectrophotometric identification does not allow the detection of other groups of flavonoids such as isoflavones or catechins that show their main peaks below 280 nm^40,102^. With respect to betalains pigments, pink-red betacyanins were distinguished from anthocyanins due to a higher wavelength of absorption peak (532-554 nm), and yellow betaxanthins were distinguished from carotenoids by the extraction in methanol solvent and the presence of only one peak at 450-500 nm^40^. In addition, it was confirmed that the family had the presence of betalians previously described. We have not quantified concentrations of each pigment group because our methods do not allow for the identification of specific compounds (i.e. type of anthocyanins, carotenoids, UAPs, etc.) present in each species. In species from Spain with whitish, cream, pale-yellow, and yellow extracts, in which we observe an absorption peak around 350-450 nm, we performed two additional tests to corroborate the presence of aurones-chalcones vs. carotenoids: color reaction in methanolic HCL extracts and differential separation with water and dichloromethane^37^.

We found some species that showed absorbance spectra peaks that did not fit any of the major pigments previously mentioned. We performed a bibliographic search to find out if there were biochemical data on the floral pigment of these species. Quinones are a rare group of pigments that may be found in some flowers, mainly anthraquinones or quinochalcones^104^. In our study, *Dipcadi serotinum* presented a compound with only one peak at 460 nm that was extracted in both methanol and acetone solutions; this was congruent with a quinone^40^, yet yellow anthraquinones have been found in other species of Asparagaceae^104^. Similarly, the compound peaking at 500 nm in all species of *Xyris* was also congruent with previously described anthraquinones for this genus^105^. To calculate the frequency of the main types of pigments, quinones were included in the anthocyanin group since they share the early steps of the biosynthetic pathway^104^. Xanthones is a rare group of flavonoids in flowers that show absorbance peaks similar to isoflavones, flavones, and flavonols^40,104^. In flowers of *Iris* spp. and *Hypericum* spp., xanthones have been previously described^106,107^, thus, they may be present, along with other UAPs, in our methanol extracts. Finally, *Adonis macrocarpa* showed a compound with a single peak at 480 nm appeared in both methanol and acetone solutions, which agreed with the red carotenoid astaxanthin previously reported in *A. aestivalis*^108^.

In species with observed UV or visible color patterns (see UAPs location in petals section), separate samples were analyzed (e.g., different floral pieces of Orchidaceae spp., apical and basal portions of ligulate/ray flowers of Asteraceae). In the Fabaceae family, we separately analyzed the banner, keels, and wings, except for species with very tiny flowers (e.g., *Trifolium* spp. or *Medicago* spp.). In these species, the total number of pigments present in a flower was the sum of the pigments found in all the different samples of a flower. The same approach was applied to species with blushing (i.e., flower color change) and flower color polymorphic species.

### UAPs location in petals

We investigated whether the presence of UAPs was located in the whole petals or produced spatial color patterns or floral guides by using UV photography^38^ and/or measuring reflectance spectra in different petal areas^34^. We considered color patterns as a spatial variation in UV or visible color within a flower, which includes UV or visible bullseyes, veins, rays, spots and/or differently colored petals/tepals (e.g., flowers of the Fabaceae or Orchidaceae; see Supplementary Data 1). These patterns have been traditionally considered as floral or nectar guides (e.g.^31,60^). The patterns were confirmed, when possible, by reviewing previous studies^3,61^.

### Floral pigments frequences in different light environments and pollination systems

In California and Spain, the studied species grow in a variety of habitats with different light environments and solar UV radiation, ranging from very low (e.g. redwood forest, oak forest, riparian) to extremely high (e.g. grasslands, desert, coastal dunes). For habitat assessments in the California Floristic Province, we used the "ecology" descriptions provided in the Jepson eFlora (www.ucjeps.berkeley.edu/eflora) and categorized them into nine main habitat types: riparian, wetland, coastal, desert, forest, grassland, woodland, montane, and rocky. When a species has more than one habitat, we chose the main habitat according to the description in Jepson eFlora, our own habitat knowledge of the species and photos of the species consulted in iNaturalist (www.inaturalist.org). In Spain, we followed the same procedure using Flora Iberica (www.floraiberica.es), categorizing species into the same habitat types as in California, with the exception of the montane habitat. For comparisons, we grouped these habitats in “shaded” and “exposed” light environments^32^, considering forest and riparian as “shaded” light environment and the rest of habitats as “exposed”.

Woodland species was not included in the analysis (N = 42 in California and 33 in S Spain) because most of the species grow in both shaded and exposed light environments. In Brazil, species grown on rocky and savanna habitats, and both are considered open habitats^93^, making comparisons between different light environments meaningless.

We categorized species from California and Brazil according to their main functional group of pollinators, i.e. “insect”, “hummingbird”, or “mixed”^88^. To assign these categories, a bibliographic search of the pollinators of all the species was carried out. (see Supplementary Table 1). Only four species from California were found with a mixed pollination syndrome and therefore were not taken into account in the analysis. In Brazil, one species showed pollination by bat and was not considered either in the analysis.

Variation in pigment frequencies between shaded and exposed environments in and between insect- and hummingbird-pollinated flowers, we studied in the three major pigment groups anthocyanins, carotenoids and chlorophylls. UAPs were not considered due to virtually all the species had these pigments.

### Statistical analysis

We performed permutation tests to know whether the number of species producing each type of flower pigment varied among the three study sites. The observed number of species having a pigment type in each study site were compared with the distribution of permuted data (1000 iterations) using the whole dataset with the three regions^109^ (Berry et al. 2019).

T-tests were used to determine whether the observed number of species showing a pigment type differs from the randomized number using two-sided p-values (α = 0.05). The same statistical analysis was used to know whether the number of pigments produced per flower varied among the study sites and to know if the frequency of anthocyanins, carotenoids and chlorophylls varies between light environments and between pollination systems. Statistical analyses were performed in R, version 4.3.2^110^ using R-studio interface. The bar graphs were created using datawrapper software (www.datawrapper.de).

## Data availability

The list of species used in this study, along with their assigned light environment and pollination system, is provided in Supplementary Data 1. Pigment data are available from the corresponding author upon reasonable request.

## Acknowledgements

We thank Elaine Meslow, Conso Barciela, Pilar Fernandez-Díaz and Julia Fernandez-Boraita for technical support, and the General Herbarium of the Universidad de Sevilla (CITIUS) for its logistical support. We sincerely thank Stacey Smith and two anonymous reviewers for their insightful comments on an earlier version of this manuscript. This study was supported by the project PID2020-116222GB-I00 funding by the Spanish government MICIU/AEI/ 10.13039/501100011033. We also want to thank grants of the Andalusian Regional Ministry of Economy, Knowledge, Business and University (PREDOC-00336 and PAIDI BIO-305, Spain), the São Paulo Research Foundation (FAPESP, Brazil) (grants #2013/50155-0, #2010/51307-0, #2009/54208-6; #2021/10639-5), the National Council for Scientific and Technological Development (CNPq, Brazil) (grants #400717/2013-1 and 306563/2022-3), and the Coordination for the Improvement of Higher Education Personnel (CAPES, Brazil) (Finance code 1 and CAPES-Print 88887.374156/2019-00). MGGC received CNPq-PDJ (#161293/2015-8) and FAPESP scholarships (#2015/10754-8, #2018/21646-0). JBW received the Santa Clara University’s WAVE grant (California, USA), EM received a REAL award (California, USA), and VR received an undergraduate research scholarship from the Northern California Botanists and TriBeta.

## Author contributions

EN, MA, PLO, MLB, LPCM and JBW designed the research; all authors collected samples; EN, JBW, MLO, MLB, IP, MGGC, NRC, VR, KC, JHM, and EM performed the biochemical analysis; EN, JCDV, MLO and MGGC analyzed the data and prepared the figures; EN, MA, JCDV, JBW, and PLO wrote the paper. All authors discussed the data and contributed to the final manuscript.

## References

1. Tong ZY, Wu LY, Feng HH, Zhang M, Armbruster WS, Renner SS, Huang SQ. 2023. New calculations indicate that 90% of flowering plant species are animal-pollinated. National Science Review 10: nwad219.

2. Frachon L, Stirling SA, Schiestl FP, Dudareva N. 2021. Combining biotechnology and evolution for understanding the mechanisms of pollinator attraction. Current Opinion in Biotechnology 70: 213–219.

3. Kay QON, Daoud HS, Stirton, C. H. 1981. Pigment distribution, light reflection and cell structure in petals. Botanical journal of the Linnean Society 83: 57–83.

4. van der Kooi CJ, Elzenga JTM, Staal M, Stavenga DG. 2016. How to colour a flower: on the optical principles of flower coloration. Proceedings of the Royal Society B: Biological Sciences 283: 20160429.

5. Caruso, C.M., Eisen, K.E., Martin, R.A. and Sletvold, N., 2019. A meta-analysis of the agents of selection on floral traits. Evolution 73: 4–14.

6. Sobel, J.M. and Streisfeld, M.A. 2013. Flower color as a model system for studies of plant evo-devo. Frontiers in Plant Science 4: 321.

7. Strauss SY, Whittall JB. 2006. Non-pollinator agents of selection on floral traits. In: Harder LD, Barret HSC, eds. Ecology and evolution of flowers. Oxford, UK: Oxford University Press, 120–38.

8. Narbona E, Arista M, Whittall JB, Camargo MGG, Shrestha M. 2021a. The role of flower color in angiosperm evolution. Frontiers in Plant Science 12: 736998.

9. Muhlemann JK, Younts TL, Muday GK. 2018. Flavonols control pollen tube growth and integrity by regulating ROS homeostasis during high-temperature stress. Proceedings of the National Academy of Sciences 115: E11188–1197.

10. Borghi M, Perez de Souza L, Yoshida T, Fernie AR. 2019. Flowers and climate change: a metabolic perspective. New Phytologist 224: 1425–1441.

11. Koski MH, Finnell LM, Leonard E, Tharayil N. 2022. Elevational divergence in pigmentation plasticity is associated with selection and pigment biochemistry. Evolution 76: 512–527.

12. Davies KM, Albert NW, Zhou Y, Schwinn KE. 2018. Functions of flavonoid and betalain pigments in abiotic stress tolerance in plants. In: Roberts J, Ed. Annual Plant Reviews Online, John Wiley & Sons Ltd., 1–41.

13. Davies KM, Landi M, van Klink JW, Schwinn KE, Brummell DA, Albert NW, Chagné D, Jibran R, Kulshrestha S, Zhou Y, Bowman JL. 2022. Evolution and function of red pigmentation in land plants. Annals of Botany 130: 613–636.

14. Pérez-Gálvez A, Viera I, Roca M. 2020. Carotenoids and chlorophylls as antioxidants. Antioxidants 9: 505.

15. Landi M, Agati G, Fini A, Guidi L, Sebastiani F, Tattini M, 2021. Unveiling the shade nature of cyanic leaves: A view from the “blue absorbing side” of anthocyanins. Plant, Cell and Environment 44: 1119–1129.

16. Ferreyra MLF, Serra P, Casati P. 2021. Recent advances on the roles of flavonoids as plant protective molecules after UV and high light exposure. Physiologia Plantarum 173: 736–749.

17. Grünig, N., Horz, J.M., Pucker, B. 2024. Diversity and ecological functions of anthocyanins. Preprint at Preprints.org, 10.20944/preprints202408.2272.v1

18. Stelzner J, Roemhild R, Garibay-Hernández A, Harbaum-Piayda B, Mock HP, Bilger W. 2019. Hydroxycinnamic acids in sunflower leaves serve as UV-A screening pigments. Photochemical and Photobiological Sciences 18: 1649–1659.

19. Nascimento LBDS, Tattini M 2022. Beyond photoprotection: The multifarious roles of flavonoids in plant terrestrialization. International Journal of Molecular Sciences 23: 5284.

20. Yonekura-Sakakibara K, Higashi Y, Nakabayashi R. 2019. The origin and evolution of plant flavonoid metabolism. Frontiers in Plant Science 10: 943.

21. Guldberg, L.D., Atsatt, P.R. 1975. Frequency of reflection and absorption of ultraviolet light in flowering plants. American Midland Naturalist 93: 35–43.

22. Liu Y, Watanabe M, Yasukawa S, Kawamura Y, Aneklaphakij C, Fernie AR, Tohge T. 2021. Cross-species metabolic profiling of floral specialized metabolism facilitates understanding of evolutional aspects of metabolism among Brassicaceae species. Frontiers in Plant Science 12: 640141.

23. Wheeler LC, Dunbar-Wallis A, Schutz K, Smith SD. 2023. Evolutionary walks through flower color space driven by gene expression in *Petunia* and allies (Petunieae). Proceedings of the Royal Society B: Biological Sciences 290: 20230275.

24. Warren J, Mackenzie S. 2001. Why are all colour combinations not equally represented as flower-colour polymorphisms? New Phytologist 151: 237–241.

25. Dalrymple RL, Kemp DJ, Flores-Moreno H, Laffan SW, White TE, Hemmings FA, Moles AT. 2020. Macroecological patterns in flower colour are shaped by both biotic and abiotic factors. New Phytologist 228: 1972–1985.

26. Dellinger, A., Maier, L., Smith, S. and Sinnott-Armstrong, M. 2024. Does the abiotic environment influence the distribution of flower and fruit colors?. Preprint at Authorea, 10.22541/au.172797328.86857709/v1.

27. Narbona E, del Valle C, Arista M, Buide ML, Ortiz PL. 2021b. Major flower pigments originate different colour signals to pollinators. Frontiers in Ecology and Evolution 9: 743850.

28. Zhang Z, Xu C, Zhang S, Shi C, Cheng H, Liu H, Zhong B. 2022. Origin and adaptive evolution of UV RESISTANCE LOCUS 8-mediated signaling during plant terrestrialization. Plant Physiology 188: 332–346.

29. Koski MH, MacQueen D, Ashman TL. 2020. Floral pigmentation has responded rapidly to global change in ozone and temperature. Current Biology 22: 4425–4431.

30. Harborne JB. 1994. Introduction to ecological biochemistry. 4th edition. Academic Press. New York. 318pp

31. Lunau K, Scaccabarozzi D, Willing L, Dixon K. 2021. A bee’s eye view of remarkable floral colour patterns in the Southwest Australian biodiversity hotspot revealed by false colour photography. Annals of Botany 128: 821–824.

32. Endler JA. 1993. The color of light in forests and its implications. Ecological Monographs 63: 1–27.

33. Shrestha M, Dyer AG, Boyd-Gerny S, Wong BB, Burd M. 2013. Shades of red: bird-pollinated flowers target the specific colour discrimination abilities of avian vision. New Phytologist 198: 301–310.

34. de Camargo MGG, Lunau K, Batalha MA, Brings S, de Brito VLG, Morellato LPC. 2019. How flower colour signals allure bees and hummingbirds: a community-level test of the bee avoidance hypothesis. New Phytologist 222: 1112–1122.

35. Rodríguez-Sambruno C, Narbona E, del Valle JC, Valido A. 2024. Bird-flower colour on islands supports the bee-avoidance hypothesis. Functional Ecology 38: 600–611.

36. Ng J, Smith SD. 2016 How to make a red flower: the combinatorial effect of pigments. AoB Plants 8: plw013.

37. Stanley LE, Ding B, Sun W, Mou F, Hill C, Chen S, Yuan YW. 2020. A tetratricopeptide repeat protein regulates carotenoid biosynthesis and chromoplast development in monkeyflowers (*Mimulus*). The Plant Cell 32: 1536–1555.

38. León-Osper M, Rossi V, Conrad K, Hernandez-Mena J, Meslow E, Fuller A, Narbona E, Whittall JW. 2024. California red hummingbird flowers: color convergence across four biochemical categories. Madroño (2^nd^ revision).

39. Mabry T, Markham KR, Thomas MB. 1970. The systematic identification of flavonoids. New York, NY, USA: Springer Science & Business Media.

40. Harborne JB. 1984. Phytochemical methods – a guide to modern techniques of plant analysis, 2nd edition. London, UK: Chapman & Hall.

41. Saha S, Singh J, Paul A, Sarkar R, Khan Z, Banerjee K. 2020. Anthocyanin profiling using UV-Vis spectroscopy and liquid chromatography mass spectrometry. Journal of AOAC International 103: 23–39.

42. Andersen ØM, Jordheim M. 2010. Chemistry of flavonoid-based colors in plants. In: Mander L, Liu H-W, eds. Comprehensive Natural Products II: Chemistry and Biology, vol. 3. Oxford, UK: Elsevier Science, 547–614.

43. Iwashina T. 2015. Contribution to flower colors of flavonoids including anthocyanins: A review. Natural Product Communications 10: 529–544.

44. Timoneda A, Feng T, Sheehan H, Walker-Hale N, Pucker B, Lopez-Nieves S, Gou R, Brockinborgton S. 2019. The evolution of betalain biosynthesis in Caryophyllales. New Phytologist 224: 71–85.

45. Verdaguer D, Jansen MA, Llorens L, Morales LO, Neugart S. 2017. UV-A radiation effects on higher plants: Exploring the known unknown. Plant Science 255: 72–81.

46. Wang M, Leng C, Zhu Y, Wang P, Gu Z, Yang R. 2022. UV-B treatment enhances phenolic acids accumulation and antioxidant capacity of barley seedlings. LWT 153: 112445.

47. Tohge, T., Wendenburg, R., Ishihara, H., Nakabayashi, R., Watanabe, M., Sulpice, R., Hoefgen, R., Takayama, H., Saito, K., Stitt, M. and Fernie, A.R., 2016. Characterization of a recently evolved flavonol-phenylacyltransferase gene provides signatures of natural light selection in Brassicaceae. Nature Communications 7: 12399.

48. Yao X, Chu J, He X, Ma C, Han C, Shen H. 2015. The changes in quality ingredients of Qi chrysanthemum flowers treated with elevated UV-B radiation at different growth stages. Journal of Photochemistry and Photobiology B: Biology 146: 18–23.

49. Wong DC, Perkins J, Peakall R. 2022. Conserved pigment pathways underpin the dark insectiform floral structures of sexually deceptive *Chiloglottis* (Orchidaceae). Frontiers in Plant Science 13: 976283.

50. Lu C, Liu Y, Yan X, Gui A, Jiang Y, Wang P, Qiao Q, Shao Q. 2024. Multiplex approach of metabolomic and transcriptomic reveals the biosynthetic mechanism of light-induced flavonoids and CGA in *Chrysanthemum*. Industrial Crops and Products 221: 119420.

51. Kaur H, Manna M, Thakur T, Gautam V, Salvi P. 2021. Imperative role of sugar signaling and transport during drought stress responses in plants. Physiologia plantarum 171: 833–848.

52. Chen W, Xiao Z, Wang Y, Wang J, Zhai R, Lin-Wang K, Espley R, Ma F, Li P. 2021. Competition between anthocyanin and kaempferol glycosides biosynthesis affects pollen tube growth and seed set of *Malus*. Horticulture Research 8: 173.

53. Bolouri-Moghaddam MR, Le Roy K, Xiang L, Rolland F, Van den Ende W. 2010. Sugar signalling and antioxidant network connections in plant cells. The FEBS journal 277: 2022–2037.

54. Zhao J. 2015. Flavonoid transport mechanisms: how to go, and with whom. Trends in Plant Science 20: 576–585.

55. Trouillas P, Sancho-García JC, De Freitas V, Gierschner J, Otyepka M, Dangles O. 2016. Stabilizing and modulating color by copigmentation: Insights from theory and experiment. Chemical Reviews 116: 4937–4982.

56. Tanaka Y, Sasaki N, Ohmiya A. 2008. Biosynthesis of plant pigments: Anthocyanins, betalains and carotenoids. Plant Journal 54: 733–749.

57. Landi M, Tattini M, Gould KS. 2015. Multiple functional roles of anthocyanins in plant–environment interactions. Environmental and Experimental Botany 119: 4–17.

58. Tohge T, Perez de Souza L, Fernie AR. 2018. On the natural diversity of phenylacylated-flavonoid and their in planta function under conditions of stress. Phytochemistry Reviews 17: 279–290.

59. Carlson, J.E. and Holsinger, K.E., 2015. Extrapolating from local ecological processes to genus-wide patterns in colour polymorphism in South African Protea. Proceedings of the Royal Society B: Biological Sciences 282: 20150583.

60. Lunau, K., De Camargo, M.G.G. and Brito, V.L.G., 2024. Pollen, anther, stamen, and androecium mimicry. Plant Biology 26: 349–368.

61. Tunes P, Camargo MGG, Guimaraes E. 2021. Floral UV features of plant species from a Neotropical savanna. Frontiers in Plant Science 12: 618028.

62. Fan XQ, Trunschke J, Ren ZX, Wang H, Pyke GH, van der Kooi CJ, Lunau K. 2024. Why are the inner and outer sides of many flower petals differently coloured? Plant Biology 26: 665–674.

63. Koski MH, Ashman TL. 2015. Floral pigmentation patterns provide an example of Gloger’s rule in plants. Nature Plants 1: 14007.

64. Todesco M, Bercovich N, Kim A, Imerovski I, Owens GL, Ruiz ÓD, Holalu SV, Madilao LL, Jahani M, Légaré JS, Blackman BK. 2022. Genetic basis and dual adaptive role of floral pigmentation in sunflowers. eLife 11: 72072.

65. Weevers TH. 1952. Flower colours and their frequency. Acta Botanica Neerlandica 1: 81–92.

66. Kevan PG. 1972. Floral colors in the high arctic with reference to insect–flower relations and pollination. Canadian Journal of Botany 50: 2289–316.

67. Dyer AG, Jentsch A, Burd M, Garcia JE, Giejsztowt J, Camargo MG, Tjørve E, Tjørve KM, White P, Shrestha M. 2021. Fragmentary blue: Resolving the rarity paradox in flower colors. Frontiers in Plant Science 15: 618203.

68. Wessinger, C.A., Rausher, M.D., 2012. Lessons from flower colour evolution on targets of selection. Journal of Experimental Botany 63: 5741–5749.

69. Kemp, J.E., Bergh, N.G., Soares, M. and Ellis, A.G., 2019. Dominant pollinators drive non-random community assembly and shared flower colour patterns in daisy communities. Annals of Botany 123: 277–288.

70. Tenhumberg, B., Dellinger, A.S., Smith, S.D., 2023. Modelling pollinator and nonpollinator selection on flower colour variation. Journal of Ecology 111: 746–760.

71. McEwen JR, Vamosi JC. 2010. Floral colour versus phylogeny in structuring subalpine flowering communities. Proceedings of the Royal Society of London B: Biological Sciences 277: 2957–2965.

72. Sargent, R.D., Ackerly, D.D., 2008. Plant–pollinator interactions and the assembly of plant communities. Trends in Ecology and Evolution 23: 123–130.

73. Albor C, Ashman TL, Stanley A, Martel C, Arceo-Gómez G. 2022. Flower colour and flowering phenology mediate plant–pollinator interaction assembly in a diverse co-flowering community. Functional Ecology 36: 2456–2468.

74. Gumbert, A., Kunze, J., Chittka, A.L., 1999. Floral colour diversity in plant communities, bee colour space and a null model. Proceedings of the Royal Society of London. Series B: Biological Sciences 266: 1711–1716.

75. Rohde, K., Papiorek, S., Lunau, K., 2013. Bumblebees (*Bombus terrestris*) and honeybees (*Apis mellifera*) prefer similar colours of higher spectral purity over trained colours. Journal of Comparative Physiology A 199: 197–210.

76. Briscoe AD, Chittka L. 2001. The evolution of color vision in insects. Annual Review in Entomology 46: 471–510.

77. Armbruster WS. 2002. Can indirect selection and genetic context contribute to trait diversification? A transition-probability study of blossom-color evolution in two genera. Journal of Evolutionary Biology 15: 468–486.

78. Irwin RE, Strauss SY, Emerson A, Guibert G. 2003. The role of herbivores in the maintenance of a flower-color polymorphism in wild radish. Ecology 84: 1733–1743.

79. Grossenbacher, D., Makler, L., McCarthy, M., Fraga, N., 2021. Abiotic environment predicts micro-but not macroevolutionary patterns of flower color in monkeyflowers (Phrymaceae). Frontiers in Plant Science 12: 636133.

80. Li, R., Zeng, Q., Zhang, X., Jing, J., Ge, X., Zhao, L., Yi, B., Tu, J., Fu, T., Wen, J., Shen, J., 2023. Xanthophyll esterases in association with fibrillins control the stable storage of carotenoids in yellow flowers of rapeseed (*Brassica juncea*). New Phytologist 240: 285–301.

81. Yuan Y, Li X, Yao X, Fu X, Cheng J, Shan H, Yin X, Kong H. 2023. Mechanisms underlying the formation of complex color patterns on *Nigella orientalis* (Ranunculaceae) petals. New Phytologist 237: 2450–2466.

82. del Valle JC, León-Osper M, Domínguez-González C, Buide ML, Arista M, Ortiz PL, Whittall JB, Narbona E. 2024. Green flowers need yellow to get noticed in a green world. Annals of Botany (doi: 10.1093/aob/mcae213).

83. Moldenke AR. 1976. California pollination ecology and vegetation types. Phytologia 34: 305–361.

84. Oliveira PE, Gibbs PE. 2000. Reproductive biology of woody plants in a Cerrado community of Central Brazil. Flora 195: 311–329.

85. Pauw A. 2019. A bird’s-eye view of pollination: Biotic interactions as drivers of adaptation and community change. Annual Review of Ecology, Evolution and Systematics 50: 477–502.

86. Gould SJ, Vrba, ES. 1982. Exaptation—a missing term in the science of form. Paleobiology 8: 4–15.

87. Brunetti, C., Sebastiani, F., Tattini, M., 2019. ABA, flavonols, and the evolvability of land plants. Plant Science 280: 448–454.

88. Dellinger, A.S., Hamilton, A.M., Wessinger, C.A., Smith, S.D., 2023. Opposing patterns of altitude-driven pollinator turnover in the tropical and temperate Americas. The American Naturalist 202: 152–165.

89. Marston A, Hostettmann K. 2006. Separation and quantification of flavonoids. In: Andersen OM, Markham KR, eds. Flavonoids: Chemistry, biochemistry and applications. Boca Raton, FL, USA: Taylor & Francis CRC Press, 1–36.

90. Davies, K.M. 2004a. An introduction to plant pigments in biology and commerce. In: Davies KM, ed. Plant pigments and their manipulation. Annual Plant Review. Oxford, UK: Blackwell Publishing Ltd, 1–22.

91. Grotewold E. 2006. The genetics and biochemistry of floral pigments. Annual Review of Plant Biology 57: 761–780.

92. Thompson JD. 2005. Plant evolution in the Mediterranean. Oxford, UK: Oxford University Press.

93. Le Stradic S, Buisson E, Fernandes GW, Morellato LPC. 2018. Reproductive phenology of two contrasting Neotropical mountain grasslands. Journal of Vegetation Sciences 29: 15–24.

94. Grant, K.A., 1966. A hypothesis concerning the prevalence of red coloration in California hummingbird flowers. The American Naturalist 100: 85–97.

95. Dafni A, Lehrer M, Kevan PG. 1997. Spatial flower parameters and insect spatial vision. Biological Reviews 72: 239–282.

96. Andersen ØM, Francis GW. 2004. Techniques of pigment identification. In: Davies KM, ed. Plant pigments and their manipulation. Annual Plant Review. Oxford, UK: Blackwell Publishing Ltd, 293–341.

97. Schoefs B. 2004. Determination of pigments in vegetables. Journal of Chromatography A 1054: 217–226.

98. Julkunen-Tiitto R, Nenadis N, Neugart S, Robson M, Agati G, Vepsäläinen J, Zipoli G, Nybakken L, Winkler B, Jansen MAK. 2015. Assessing the response of plant flavonoids to UV radiation: an overview of appropriate techniques. Phytochemistry Reviews 14: 273–297.

99. del Valle JC, Gallardo-López A, Buide ML, Whittall JB, Narbona E. 2018. Digital photography provides a fast, reliable, and noninvasive method to estimate anthocyanin pigment concentration in reproductive and vegetative plant tissues. Ecology and Evolution 8: 3064–3076.

100. Lee J, Rennaker C, Wrolstad RE. 2008. Correlation of two anthocyanin quantification methods: HPLC and spectrophotometric methods. Food Chemistry 110: 782–786.

101. Thrane J-E, Kyle M, Striebel M, Haande S, Grung M, Rohrlack T, Andersen T. 2015. Spectrophotometric analysis of pigments: A critical assessment of a high-throughput method for analysis of algal pigment mixtures by spectral deconvolution. PLoS ONE 10: e0137645.

102. Ritchie RJ. 2006. Consistent sets of spectrophotometric chlorophyll equations for acetone, methanol and ethanol solvents. Photosynthesis Research 89: 27–41.

103. Fossen T, Andersen ØM. 2006. Spectroscopic techniques applied to flavonoids. In: Andersen OM, Markham KR, eds. Flavonoids: Chemistry, biochemistry and applications. Boca Raton, FL, USA: Taylor & Francis CRC Press, 37–142.

104. Davies KM. 2004b. Important rare plant pigments. In: Davies KM, ed. Plant pigments and their manipulation. Annual Plant Review. Oxford, UK: Blackwell Publishing Ltd, 214–247.

105. Cota BB, de Oliveira AB, Guimarães KG, Mendonça MP, de Souza Filho JD, Braga FC. 2004. Chemistry and antifungal activity of *Xyris* species (Xyridaceae): a new anthraquinone from *Xyris pilosa*. Biochemical Systematics and Ecology 32: 391–397.

106. Iwashina T, Mizuno T. 2020. Flavonoids and xanthones from the genus Iris: Phytochemistry, relationships with flower colors and taxonomy, and activities and function. Natural Product Communications 15: 1–35.

107. Zeliou K, Koui EM, Papaioannou C, Koulakiotis NS, Iatrou G, Tsarbopoulos A, Papasotiropoulos V, Lamari FN. 2020. Metabolomic fingerprinting and genetic discrimination of four Hypericum taxa from Greece. Phytochemistry 174: 112290.

108. Cunningham FX, Gantt E. 2011. Elucidation of the pathway to astaxanthin in the flowers of *Adonis aestivalis*. The Plant Cell 23: 3055–3069.

109. Berry KJ, Johnston JE, Mielke PW. 2019. A primer of permutation statistical methods. Springer International Publishing.

110. R. Core Team, 2024. R: A language and environment for statistical computing.

