## Supplementary Table 1 for "Transcontinental patterns in floral pigment frequencies among animal-pollinated species"

**Supplementary Table 1.** The effect of light environment (exposed vs. shaded) on major pigment distribution in California and S Spain. T-tests were used to determine whether the observed number of species showing a pigment type differs from the randomised number using 1000 random permutations. Significant two-sided p-values are marked with an asterisk.

| **Region** | **Pigment** | **Observed N**  **(exposed/shaded)** | **Expected N**  **(exposed/shaded)** | **P-value** |
| --- | --- | --- | --- | --- |
| California |  |  |  |  |
|  | Anthocyanins | 187/35 | 184/38 | 0.150 |
|  | Carotenoids | 145/25 | 138/31 | 0.120 |
|  | Chlorophylls | 60/19 | 65/14 | 0.084 |
| S Spain |  |  |  |  |
|  | Anthocyanins | 166/16 | 162/17 | 0.094 |
|  | Carotenoids | 110/12 | 108/14 | 0.444 |
|  | Chlorophylls | 48/15 | 56/7 | 0.002* |
