## Supplementary Table 2 for "Transcontinental patterns in floral pigment frequencies among animal-pollinated species"

**Supplementary Table 2.** The effect of pollination system (insect- vs. hummingbird-pollination) on major pigment distribution in California and SE Brazil. T-tests were used to determine whether the observed number of species showing a pigment type differs from the randomised number using 1000 random permutations. Significant two-sided p-values are marked with an asterisk. Species with mixed pollination system were not considered due to the low sample size (see Star Methods).

| **Region** | **Pigment** | **Observed N**  **(insect/hummingbird)** | **Expected N**  **(insect/hummingbird)** | **P-value** |
| --- | --- | --- | --- | --- |
| California |  |  |  |  |
|  | Anthocyanins | 199/45 | 219/25 | <0.001* |
|  | Carotenoids | 153/28 | 164/17 | 0.004* |
|  | Chlorophylls | 89/3 | 82/10 | 0.006* |
| SE Brazil |  |  |  |  |
|  | Anthocyanins | 55/6 | 56/5 | 0.188 |
|  | Carotenoids | 32/4 | 33/3 | 0.202 |
|  | Chlorophylls | 10/1 | 10/1 | 0.432 |
