## Supplementary Table 3 for "Transcontinental patterns in floral pigment frequencies among animal-pollinated species"

**Supplementary Table 3.** Example compounds for each main pigment group used in Figure 1.

| **Main pigment group** | **Example compound** | **Reference** |
| --- | --- | --- |
| Hydroxycinnamates | *p*-coumaric acid | Miyagusuku-Cruzado et al., 2022 |
|  | Caffeic acid | Miyagusuku-Cruzado et al., 2022 |
| Flavones | Luteolin | Mabry et al., 1970 |
| Flavonols | Quercetin 3,7-O-diglucoside | Mabry et al., 1970 |
| Chalcones | 2,2',4-trihydroxychalcone | Mabry et al., 1970 |
| Aurones | Aaritimetin | Mabry et al., 1970 |
| Pelargonidins | Pelargonidin-3-0-rutinoside | Qin et al., 2010 |
| Cyanidins | Cyanidin-3-0-glucuside | Qin et al., 2010 |
| Delphinidis | Delphinidin-3-0-rutinoside | Merken & Beecher, 2000 |
| Betaxanthins | Amaranthine | Cai et al., 2001 |
| Betacyanins | Miraxanthin | Cai et al., 2001 |
| Luetins | Luetin | Zang et al., 1997 |
| Zeaxanthins | Zeaxanthins | Zang et al., 1997 |
| Astaxanthins | Adonixanthin | Cunningham & Gantt, 2011 |
| Chlorophyll *a* | Chlorophyll *a* | Sawicki et al., 2019 |
| Chlorophyll *b* | Chlorophyll *b* | Sawicki et al., 2019 |
